# Improving AlphaFold2 and 3-based protein complex structure prediction with MULTICOM4 in CASP16

**DOI:** 10.1101/2025.03.06.641913

**Authors:** Jian Liu, Pawan Neupane, Jianlin Cheng

**Affiliations:** Department of Electrical Engineering & Computer Science, NextGen Precision Health, University of Missouri, Columbia, Missouri, 65211, United States of America

**Keywords:** KEYWORD S: Protein structure prediction, protein model quality assessment, deep learning, artificial intelligence

## Abstract

With AlphaFold achieving high-accuracy tertiary structure prediction for most single-chain proteins (monomers), the next major challenge in protein structure prediction is accurately modeling multi-chain protein complexes (multimers). We developed MULTICOM4, the latest version of the MULTICOM system, to improve protein complex structure prediction by integrating transformer-based AlphaFold2, diffusion model-based AlphaFold3, and our in-house techniques. These include protein complex stoichiometry prediction, diverse multiple sequence alignment (MSA) generation leveraging both sequence and structure comparison, modeling exception handling, and deep learning-based model quality assessment.

MULTICOM4was blindly evaluated in the 16th communitywide Critical Assessment of Techniques for Protein Structure Prediction (CASP16) in 2024. In Phase 0 of CASP16, where stoichiometry information was unavailable, MULTICOM predictors performed best, with MULTICOM_human achieving a TM-score of 0.752 and a DockQ score of 0.584 for top-ranked predictions on average. In Phase 1 of CASP16, with stoichiometry information provided, MULTICOM_human remained among the top predictors, attaining a TM-score of 0.797 and a DockQ score of 0.558 on average. The CASP16 results demonstrate that integrating complementary AlphaFold2 and 3 with enhanced MSA inputs, comprehensive model ranking, exception handling, and accurate stoichiometry prediction can effectively improve protein complex structure prediction.

## 1 INTRODUCTION

Accurate prediction of quaternary structures of protein complexes is crucial for understanding protein-protein interaction and protein function. In recent years, deep learning-based methods such as AlphaFold2-Multimer[1, 2] and AlphaFold3[3] have significantly advanced quaternary structure modeling. However, the accuracy of protein complex structure prediction is still much lower than that of protein tertiary structure prediction. Significant challenges remain in handling large protein complexes (assemblies), predicting structures for complexes with shallow or non-informative MSAs (e.g., antibody-antigen complexes), and ranking structural models accurately. Additionally, the lack of accurate methods for protein complex stoichiometry prediction prevents applying AlphaFold2 and 3 to many protein complexes whose stoichiometry (subunit count) is still unknown.

To address these challenges, we developed MULTICOM4, the latest version of our MULTICOM protein structure prediction system [4, 5, 6], on top of AlphaFold2, AlphaFold3, and our in-house modeling techniques. Because AlphaFold2 and AlphaFold3 use two different deep learning architectures to generate complex structural models - a transformer-based architecture and a diffusion model-based architecture, they have complementary strengths, even though AlphaFold3 is somewhat more accurate than AlphaFold2 on average [3]. MULTICOM4 uses both AlphaFold2 and AlphaFold3 to generate structural models.

MULTICOM4 enhances AlphaFold-based structural model generation by feeding it with diverse multimeric MSAs generated by both sequence and structure alignment tools. It systematically leverages species information, protein-protein interaction, and structural similarity to improve MSA construction for protein complexes. The diverse MSAs containing co-evolutionary signals [4] are used by AlphaFold with different modeling parameters to generate a relatively large pool of structural models for model ranking and selection.

MULTICOM4 integrates multiple model ranking scores / methods, including AlphaFold2-Multimer confidence scores, AlphaFold3 ranking score, GATE[7], VoroMQA[8], GCPNet-EMA[9], EnQA[10], and a structural consensus score (i.e., average structural TM-score between a model and other models)[11, 12], to enhance model ranking.

Moreover, to handle very large protein complexes with thousands of residues that are too big to be modeled by AlphaFold or unusual hard targets whose interaction interfaces cannot be predicted well, special modeling techniques, such as the divide and conquer modeling technique and the traditional template-based modeling technique, are used by MULTICOM4 to handle them as exceptions.

Finally, to meet the increasing need of predicting the structure for protein complexes whose stoichiometry information is unknown, MULTICOM4 applies PreStoi[13], a novel stoichiometry prediction method based on AlphaFold3 structure prediction and template information, to predict stoichiometry to guide structural model generation.

The initial development of the MULTICOM4 system based on AlphaFold2 was completed days before it participated in CASP16 that started on May 1, 2024. AlphaFold3-based model generation was added into MULTICOM4 after the AlphaFold3 web server was released to the public during the early stage of the CASP16 experiment and was used more and more as CASP16 progressed. Five different variants of MULTICOM4 were blindly tested in CASP16 as three server predictors (MULTICOM_AI, MULTICOM_GATE, MULTICOM_LLM) and two human predictors (MUL-TICOM, MULTICOM_human). Some components in these predictors, mostly model ranking methods, continued to evolve during the CASP16 experiment until they stabilized.

MULTICOM4 achieved outstanding performance in CASP16 by obtaining the best results in predicting the structures of protein complexes without stoichiometry information in Phase 0 and ranking among the top predictors in predicting the structures of protein complexes with known stoichiometry in Phase 1. It also outperformed both standard AlphaFold2 and AlphaFold3. Therefore, the special modeling techniques in MULTICOM4 can be used by the community to further improve the prediction of complex structures based on AlphaFold2 and 3 and broaden their application to the vast majority of protein complexes without stoichiometry information.

## 2 MATERIALS AND METHODS

During CASP16, multimeric targets were released in three distinct phases, each introducing unique challenges and opportunities for structure prediction: (1) Phase 0: Targets were released with sequences of subunits but unknown stoichiometry (counts of subunits). Predictors were tasked with predicting both the stoichiometry and quaternary structure. (2) Phase 1: The true stoichiometry information for the targets used in Phase 0 was revealed, requiring predictors to predict the quaternary structure based on this correct stoichiometry. Some new targets not used in Phase 0 were also released in Phase 1. (3) Phase 2: Predictions submitted by CASP16 participants during Phase 0 and Phase 1, along with a large number of structures generated by MassiveFold[14], were released. Predictors were asked to predict the quaternary structures of the same targets used in Phase 1 and could leverage the Phase 1 predictions of other CASP16 predictors. The prediction methods of MULTICOM4 used in the three phases are described below.

### 2.1 Stoichiometry Prediction

For protein complexes, whose stoichiometry information is not available, MULTICOM4 applies PreStoi[13] that integrates AlphaFold3 and template information to predict the stoichiometry for the targets. Specifically, it uses the structural templates in the Protein Data Bank (PDB)[15] found for each subunit of a protein complex to set a range for the number of copies of the subunit. The ranges are used to propose a small number of candidate stoichiometries for the complex. AlphaFold3 is then used to generate dozens of structural models for each candidate. The maximum and average AlphaFold3 ranking scores of the models of the candidates are used to rank them. Moreover, if a template in the PDB that can cover all the subunits of the complex can be found, the stoichiometry of the complex can be directly inferred from that of the template and is used as the preferred stoichiometry. Usually, one to three stochiometries were predicted for CASP16 Phase 0 targets, depending on their difficulty. The stoichiometry prediction was only applied to the Phase 0 targets in CASP16.

### 2.2 Model Generation

The model generation and ranking process of MULTICOM4 is illustrated in Figure 1. During CASP16, given a protein complex target with predicted or known stoichiometry in Phase 0 or Phase 1, MULTICOM4 used the locally installed AlphaFold2-Multimer (simply called AlphaFold2) and the AlphaFold3 web server to generate structural models for the target. The AlphaFold3 web server was used because its software was not available during the CASP16 experiment.

**FIGURE 1.**
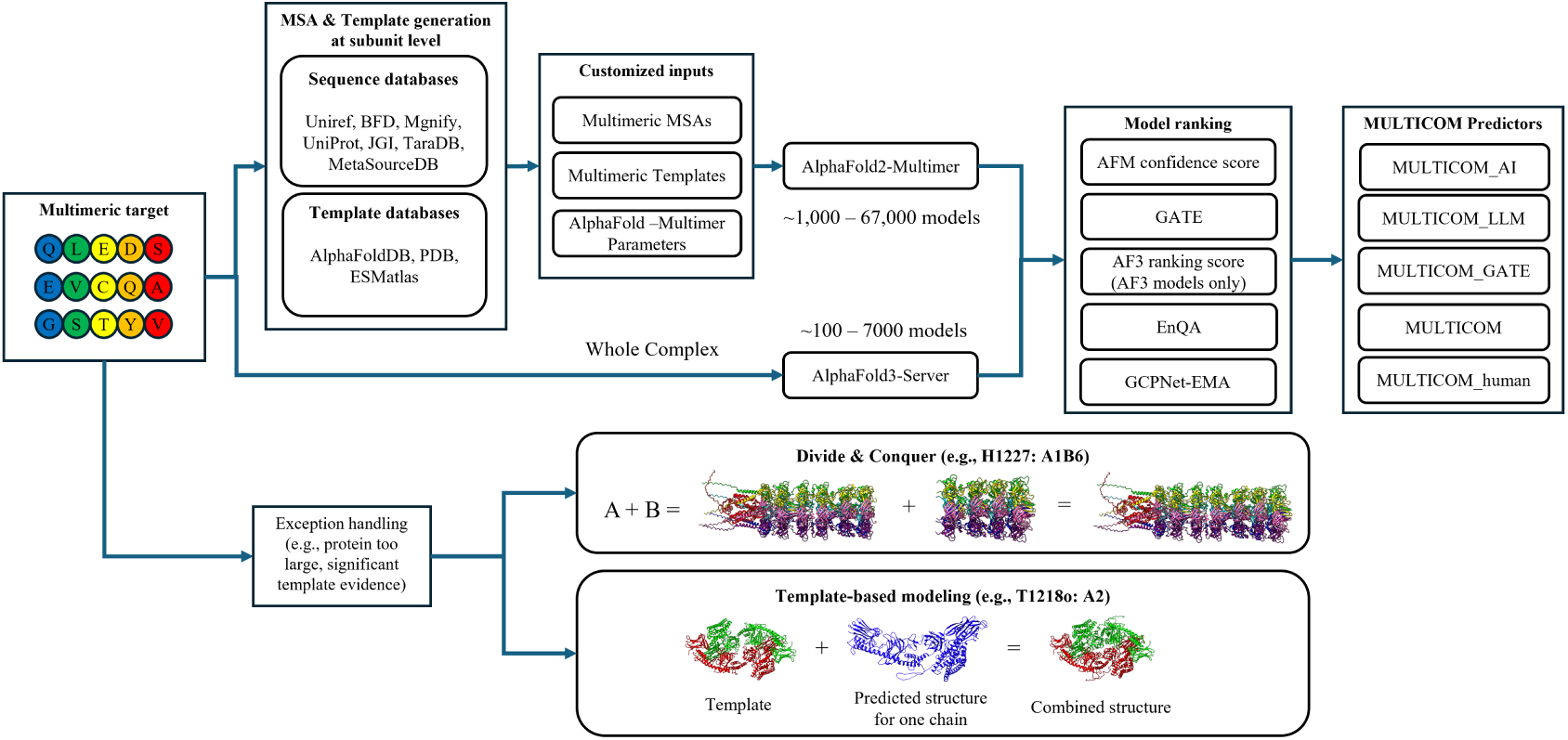
The workflow of model generation and ranking in the MULTICOM4 system.

#### 2.2.1 AlphaFold2-based Model Generation

To make AlphaFold2 more likely generate some high-quality models, MULTICOM4 uses multiple complementary techniques to generate diverse MSAs and templates for a target and use them as input for AlphaFold2 with different parameter settings to generate a pool of structural models.

##### Subunit MSA Generation and Template Identification

For each subunit in a protein complex, two types of multiple sequence alignments (MSAs), sequence alignment-based MSAs and structure alignment-based MSAs, are generated as follows.

Sequence-based MSAs are constructed by searching the sequence of each subunit against a comprehensive set of sequence databases, including UniRef30[16], UniRef90[17], Metaclust[18], BFD[19, 18], Mgnify[20], UniProt[17], JGI[21], TaraDB[22], and MetaSourceDB[23]. These searches were conducted using HHblits[24], JackHmmer[25], and DeepMSA2[26] to construct MSAs that capture sequence co-evolutionary signals within the subunit.

Structure-based MSAs are generated by first predicting 850–1,900 tertiary structural models for each subunit using AlphaFold2[1]. The highest-ranked model, selected based on AlphaFold’s global plDDT score, is then searched against structural databases, including AlphaFoldDB[27] and the ESM Metagenomic Atlas[28], using FoldSeek[29]. The FoldSeek structural alignments between the subunit and the similar structural hits are used to construct a structure-based MSA for the subunit, complementing the sequence-based MSAs.

Structural templates for each subunit of a protein complex are separately identified from PDB70, pdb_complex[5], and pdb_sort90[6] using HHsearch[30]. These templates can be used as input for AlphaFold2 to enhance structural accuracy by incorporating experimentally determined structural information.

##### Complex MSA Construction

To construct MSAs for a complex, subunit sequence-based MSAs are concatenated using species information[2, 31], UniProt accession IDs, STRING interactions[32], and PDB protein complexes. This approach generated 13 types of concatenated MSAs for heteromers and 7 types for homomers (excluding STRING interactions and UniProt accession IDs for homomers). Additionally, the DeepMSA2 pairing protocol is applied, where MSAs generated for each subunit by DeepMSA2 are ranked based on the highest plDDT score of AlphaFold2 structural predictions using the corresponding MSA. The top-ranked MSAs are then concatenated, producing up to 20 types of multimeric MSAs, depending on the availability of MSAs and the number of subunits in the complex.

To construct structure-based MSAs for a complex, sequence alignments in the subunit structure-based MSAs that shared the same identifier are concatenated to enhance co-evolutionary information between subunits. Additionally, unpaired alignments in the subunit structure-based MSAs are also included to provide complementary structural information.

##### Complex Template Construction

Subunit templates are concatenated if they share the same PDB code as the templates for a protein complex. If the total number of concatenated templates is fewer than four, the top-ranked unpaired subunit templates are selected to ensure a minimum of four templates.

##### Modeling Parameters

To generate a diverse set of structural models, various AlphaFold2-Multimer parameters are used with MULTICOM4. Key configurations included setting num_ensemble to 1 and varying num_recycle among 3, 20, or 21[33] to introduce structural variability. The model_preset is alternated between multimer_v1, multimer_v2, and multimer_v3 to leverage different sets of pretrained weights. Additionally, the dropout setting is adjusted by enabling or disabling dropout in the Evoformer network and structural module[33], further contributing to model diversity.

##### Complex Structure Prediction

The constructed complex MSAs, complex templates, and diverse modeling parameters are used to produce up to 67 sets of inputs for AlphaFold2-Multimer to generate structural models. Depending on computational constraints and target complexity, this approach produced between 1,000 and 67,000 structural models per target during CASP16, ensuring broad structural diversity and capturing multiple plausible conformations.

#### 2.2.2 AlphaFold3-based Model Generation

During CASP16, MULTICOM4 used the AlphaFold3 web server to generate structural models for protein complexes. To ensure structural diversity, multiple AlphaFold3 prediction runs were performed per target, generating 100 to 7,000 models. Since AlphaFold3 software was released after CASP16 ended, it can be called locally by MULTICOM4 to generate structural models with customized inputs.

#### 2.2.3 Exception Handling

The MULTICOM4 system applies specialized strategies to address unique challenges posed by exceptionally large protein complexes (e.g., more than 5000 residues) and hard targets with significant structural homologs whose models generated by AlphaFold have low confidence/ranking scores. For a large complex (e.g., H1227) exceeding the length limit of AlphaFold, a divide-and-conquer approach is employed, where the protein is partitioned into smaller overlapped chunks. The structures of these chunks are predicted separately and subsequently assembled into a complete model.

For targets that have significant structural templates but whose AlphaFold-predicted structures have low confidence (e.g., T1218o), MULTICOM4 uses Modeller[34] to build models for them from the templates and then combine the template-based complex models with the structural models of their subunits built by AlphaFold. The reason of incorporating the AlphaFold models of the subunits is that AlphaFold can usually accurately predict the structures of individual subunits even when it cannot accurately predict the structure of a whole complex.

### 2.3 Model Ranking for MULTICOM Predictors in Phases 0, 1, and 2

Five variants of MULTICOM4 participated in the Phase 0 experiment in CASP16 as three server predictors (MULTI-COM_AI, MULTICOM_GATe, and MULTICOM_LLM) and two human predictors (MULTICOM and MULTICOM_human). The server predictors had three days to make predictions, while the human predictors were given two to three weeks. They used the same model generation method to generate structural models, but applied different model ranking strategies to select top 5 predicted structures as follows.

MULTICOM_AI ranked models using AlphaFold2-Multimer confidence scores. The AlphaFold2-Multimer confidence score for AlphaFold3-generated models can be calculated by the formula: 0.8 × ipTM + 0.2 × pTM. MULTI-COM_GATE utilized GATE to rank models. GATE uses a graph transformer to integrate multiple model quality scores to predict the quality for each model in a model pool.

MULTICOM_LLM used the average score of GATE and AlphaFold2-Multimer confidence scores to select models in the early stage of CASP16. Later, AlphaFold3 ranking scores were used to identify the top-1 model among AlphaFold3-generated models, while the remaining top models were selected by the average of AlphaFold2-Multimer confidence score and GATE score.

MULTICOM initially used the AlphaFold2-Multimer confidence score to select models. After AlphaFold3 was integrated into the model generation process, the top-1 model was selected based on the average of the AlphaFold3 ranking score and GATE score, while the remaining top models were chosen using additional ranking metrics (e.g., the average of GATE and AlphaFold2-Multimer confidence scores, GATE, GCPNet-EMA, EnQA, and average pairwise similarity score), with human intervention if necessary. For some large targets, if models were only generated by AlphaFold3 because AlphaFold2 could not generate (reasonable) models in a short time, the AlphaFold3 ranking scores were used to select the top-1 model.

MULTICOM_human initially used the GATE score to select top models. Later, multiple ranking metrics (e.g., AlphaFold2-Multimer confidence score, GATE, GCPNet-EMA, EnQA, average pairwise similarity score were considered, and the top-1 model was selected based on the average of GATE and AlphaFold2-Multimer confidence scores, with human oversight if necessary. After AlphaFold3 was fully incorporated into model generation, models were ranked using AlphaFold2-Multimer confidence scores with manual intervention if necessary. For AlphaFold3-only model pools, the top-1 model was chosen based on the average of the AlphaFold3 ranking score and the AlphaFold2-Multimer confidence score.

To ensure structural diversity for hard targets, while always keeping top-1 ranked model, MULTICOM_AI and MULTICOM_human selected additional 4 models based on both ranking and structural dissimilarity (TM-score with the already selected models < 0.8) and MULTICOM_GATE and MULTICOM_LLM selected additional 4 models from different model clusters generated by the K-means clustering. After AlphaFold3 was fully incorporated into the system, all MULTICOM predictors ensured that, if an AlphaFold3 model was absent from the initial top 5 selections, the lowest-ranked model in the top 5 was replaced by the highest-ranked AlphaFold3 model.

In addition to the main difference in model ranking, the five MULTICOM predictors used slightly different variants of PreStoi to predict stoichiometry in Phase 0, resulting in some differences in stoichiometry prediction and, consequently, differences in their accuracy of complex structure prediction.

The same five predictors above participated in the Phase 1 prediction. They used the same model generation and ranking strategies. For the same targets used in both Phase 0 and Phase 1, in Phase 1, they incorporated previously generated Phase 0 models with correct stoichiometry if there were any. Therefore, the number of models available to use in Phase 1 was usually much larger than in Phase 0.

Only two human MULTICOM predictors participated in Phase 2. They did not generate new in-house models. Instead, they simply ranked and selected models from all the models submitted by CASP16 predictors, 10 top-ranked MassiveFold models, and 20 in-house top-ranked AlphaFold3-generated models and refined them. Therefore, in this work, we focus on describing the MULTICOM predictors and results in Phases 0 and 1.

## 3 RESULTS

We use the following six complementary metrics to assess both the global fold accuracy, local structural accuracy, and interface accuracy of the structures predicted by the MULTICOM predictors during CASP16. Interface Quality Metrics. The ICS (Interface Contact Score)[35] and IPS (Interface Packing Score)[35] are used to evaluate inter-subunit interaction interface. The DockQ_wave[36] (a weighted average of the DockQ[37] scores on all interfaces) and the QS-best score[38] further assess the interface quality. Local Structural Quality Metrics. The lDDT (Local Distance Difference Test)[39] measured the accuracy of local residue-level predictions. Global Structural Quality Metrics. The TM-score (Template Modeling Score)[40] quantifies global structural similarity between predicted and native complex structures.

The quality scores for the structures predicted by both MULTICOM predictors and other CASP16 predictors were downloaded from the CASP16 website. We focus on evaluating the quality of the top-1 model for each target predicted by the predictors if not stated otherwise, even though the best of five models for each target may be considered in some analysis. In addition to the original metric score of a model, Z-score of the model of a predictor relative to the models predicted by other predictors is also used. The sum of Z-scores across all targets within each phase is used to compare the performance of the predictors.

### 3.1 Protein Complex Prediction Results without Stoichiometry Information (Phase 0)

#### 3.1.1 Comparison between MULTICOM Predictors and other CASP16 predictors in Phase 0

There are 26 valid complex targets released in Phase 0. Figure 2(A) illustrates the sum of the Z-scores of top-1 predictions across all the targets in terms of all six evaluation metrics for the top 20 out of 67 CASP16 predictors. Four MULTICOM predictors (MULTICOM_AI, MULTICOM_human, MULTICOM_GATE, and MULTICOM) ranked 1st to fourth, substantially outperforming all other CASP16 predictors, while MULTICOM_LLM ranked no. 7. The difference in the performance of the MULTICOM predictors was due to the difference in the stoichiometry prediction and model ranking. Overall, MULTICOM predictors consistently achieved high Z-scores on most targets in interface, local, and global structural metrics, such as ICS, IPS, DockQ, lDDT, QS-best, and TM-score, demonstrating their robustness and reliability in handling multimeric targets with unknown stoichiometry.

**FIGURE 2.**
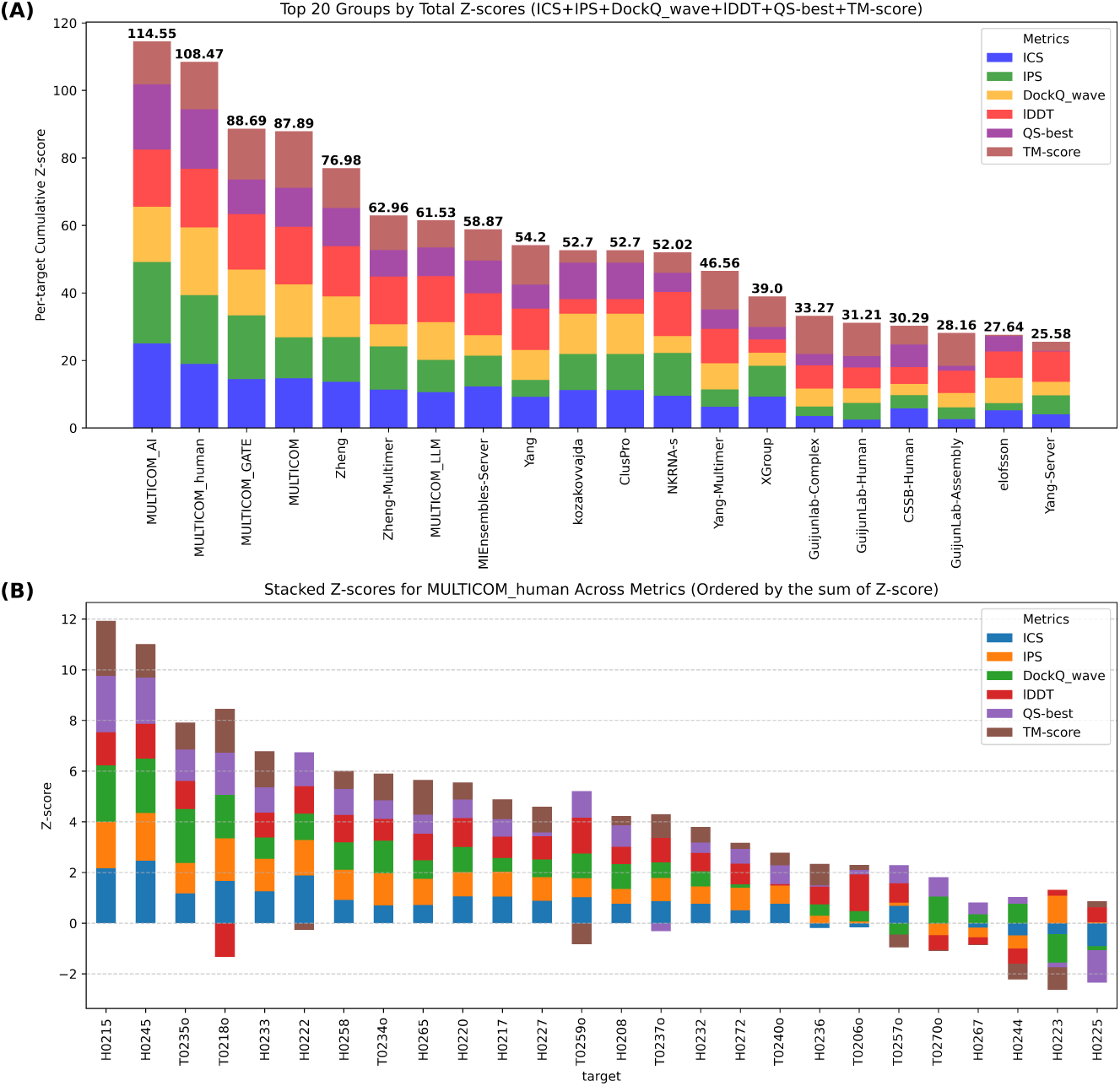
The overall performance of top 20 out of 67 CASP16 predictors and the detailed performance of MULTICOM_human across 26 Phase 0 targets. (A) The cumulative Z-scores of the six metrics for top 20 predictors on 26 targets. The standard AlphaFold3 predictor run by Elofsson’s group ranked 34th with a cumulative Z-score-46.96 (not shown); (B) The per-target Z-score of the top-1 prediction of MULTICOM_human for each target.

Here, we selected MULTICOM_human to investigate how it outperformed most other CASP16 predictors in detail because it performed well in both Phase 0 and Phase 1. Figure 2(B) illustrates per-target Z-scores of MULTI-COM_human. MULTICOM_human consistently performed above average (Z-score > 0) in terms of most evaluation metrics, particularly excelling on 20 targets with the sum of Z-scores of the six metrics higher than 2, such as H0215 (Z-score = 11.93), H0245 (Z-score = 11.01), T0235o (Z-score = 7.92), T0218o (Z-score = 7.13), and H0233 (Z-score = 6.79).

H0215 (Z-score = 11.93, Figure 3) is a nanobody target with stoichiometry A1B1 on which most CASP16 predictors failed. The top-1 model of MULTICOM_human was generated using AlphaFold2-Multimer. Several AlphaFold2-Multimer variants with different sampling parameters in MULTICOM4 produced models of near-native quality, achieving a TM-score of around 0.987 and a DockQ_wave of around 0.863. Particularly using multimer_v1 weights to generate 1,000 models produced some very high-quality models. In contrast, the top-ranked AlphaFold3-generated model has a low TM-score of 0.683 and DockQ_wave of 0.125, indicating AlphaFold3 failed to predict the correct interface between the two subunits, even though the tertiary structures of the individual subunits were well predicted. This example highlights the importance of using different modeling parameters with AlphaFold to generate better models. For H0245 (Z-score = 11.01, Figure 3), a FUNComplex target with stoichiometry A1B1, the top-1 model of MULTICOM_human was generated from the FoldSeek structure-based MSAs with AlphaFold2-Multimer. The resulting model achieved a TM-score of 0.94 and DockQ_wave of 0.808, outperforming the AlphaFold3 model, which had a TM-score of 0.84 and DockQ_wave of 0.379. This case shows that structure-based MSAs can help generate better models than sequence-based MSAs for some hard targets whose subunits have few homologous sequences.

**FIGURE 3.**
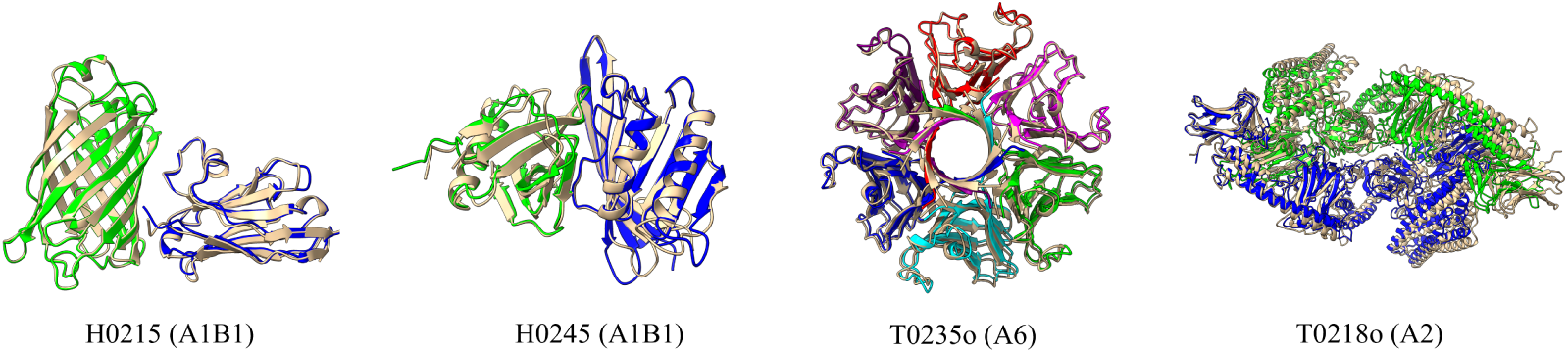
Four good examples (H0215, H0245, T0235o, T0218o) predicted by MULTICOM_human. Native structure is colored in gold, while the subunits of the predicted structure are highlighted in different colors.

For T0235o (Z-score = 7.92, Figure 3), a target with stoichiometry A6, MULTICOM_human’s advantage was partly attributed to its correct prediction of stoichiometry as A6, which was correctly predicted by only 23 out of 44 predictors that submitted predictions for this target. The top AlphaFold2-Multimer model, selected as the top-1 model of MULTICOM_human, achieved a TM-score of 0.988 and DockQ_wave of 0.794, outperforming the AlphaFold3 model with a TM-score of 0.905 and DockQ_wave of 0.278. This example shows that the correct stoichiometry prediction is critical for generating high-quality complex structures. In fact, MULTICOM_human achieved a stoichiometry pre-diction accuracy of 67.9% for top-1 models in Phase 0, higher than most other CASP16 predictors, which gave it a big advantage in Phase 0 complex structure prediction because models with correct stoichiometry are usually much more accurate than ones with incorrect stoichiometry.

For T0218o (Z-score = 7.13, Figure 3), MULTICOM_human excelled because it correctly predicted the stoichiometry of A2 from significant template hits (e.g., 4W8J, e-value = 7E-275), while most CASP16 predictors incorrectly predicted a stoichiometry of A3 probably by using AlphaFold2 or AlphaFold3 modeling to select A3, and it combined the complex structural model built from the templates with the tertiary structural models of the subunit predicted by AlphaFold3. The two identical subunits of this target have an unusual N-terminal region - C-terminal region cross interaction, which could not be correctly predicted by either AlphaFold2 or 3 because they always tried to build either N-terminal - N-terminal interaction or C-terminal - C-terminal interaction. However, the exception handling in MULTICOM4 was able to use the template-based modeling to correctly predict it. The exception handling improved the scores of five out of six metrics well above average, but caused a drop in lDDT score below average, indicating the template-based modeling substantially improved the prediction accuracy of the overall fold and the interface but caused some quality loss in local structures.

Another good exception handling example is a large complex H0227 (stoichiometry: A1B6) with 5689 residues (Figure 4), which exceeded the length limit of both AlphaFold2-Multimer and AlphaFold3 web server. MULTICOM_human divided it into two components: a component of a full subunit A and 6 partial subunit B with the first 745 residues and a component of 6 partial subunit B with residues from 390 to 877. The models generated for the two components by AlphaFold3 were combined to build full-length models via superimposing the models of the two components through their overlapped regions, yielding a high-quality top-1 model for MULTICOM_human with TM-score of 0.914 and DockQ_wave score of 0.533. Similarly, this divide and conquer strategy also enabled MULTICOM_human built a better model for two very large targets: H0217 (stoichiometry: A2B2C2D2E2F2) with 5878 residues and H0272 (stoichiometry: A1B1C1D1E1F1G1H1I1) with 6879 residues using AlphaFold3. Overall, these four examples demonstrate the importance of using exception handling to deal with the failure or limitation of AlphaFold.

**FIGURE 4.**
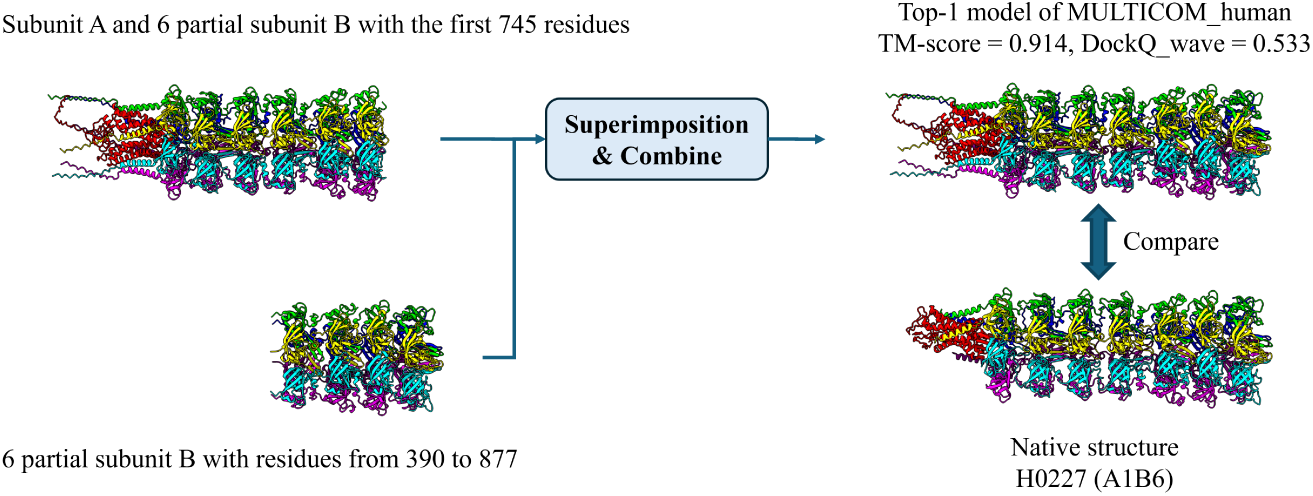
Application of the divide and conquer strategy for H0227 (stoichiometry: A1B6). The top-1 model of MULTICOM_human has TM-score of 0.914 and DockQ_wave score of 0.533.

For H0233 (Z-score = 6.79), an antibody-antigen target with stoichiometry A2B2C2, AlphaFold2-Multimer and AlphaFold3 produced significantly different structural predictions. While AlphaFold3 models were ranked using their own ranking scores, their AlphaFold2-Multimer confidence scores were still calculated for direct comparison with AlphaFold2-Multimer-generated models. Notably, AlphaFold3 models had higher confidence scores (e.g., 0.806) than AlphaFold2-Multimer models (e.g., 0.709), leading to the selection of a top AlphaFold3 model as the top-1 model for MULTICOM_human. The top-1 MULTICOM_human model achieved a TM-score of 0.991 and DockQ_wave of 0.882, much higher than TM-score of 0.493 and DockQ_wave of 0.581 of the top-ranked AlphaFold2-Multimer model. The higher structural accuracy of the AlphaFold3 model in this case demonstrates its effectiveness in predicting the structure of this antibody-antigen complex.

Despite its strong performance across most targets, MULTICOM_human performed worse than average CASP16 predictors (e.g., Z-score < 0) with certain targets. For H0244 (Z-score =-1.18), a nanobody target with stoichiometry A2B2C2, the failure stemmed from an incorrect stoichiometry prediction as A1B1C1. The top-1 model of MULTI-COM_human, based on the incorrect stoichiometry A1B1C1, has a TM-score of 0.332, while the best of its top 5 models was generated by AlphaFold3 with the correct A2B2C2 stoichiometry, achieving a TM-score of 0.671. The similar problem occurred with H0267 (Z-score =-0.05) with correct stoichiometry A2B2 (hetero tetramer). MULTI-COM_human incorrectly used stoichiometry A1B1 for its top-1 model, while including both AlphaFold2-Multimer and AlphaFold3 models with stoichiometry A2B2 within top 5 models. These two examples highlight the importance of correctly predicting stoichiometry for better complex structure prediction.

For H0223 (Z-score =-1.32), an antibody-antigen target with stoichiometry A1B1C1, conformations generated by AlphaFold2-Multimer and AlphaFold3 differed significantly, making it challenging to determine which was better. MULTICOM_human selected a sub-optimal model generated by AlphaFold2-Multimer with a TM-score of 0.763 and DockQ_wave of 0.329 as top-1 model, which was significantly worse than a good AlphaFold3 model with a TM-score of 0.941 and a DockQ_wave score of 0.535. For H0225 (Z-score =-1.47), another antibody target with stoichiometry A1B1C1, the top-1 model of MULTICOM_human was generated by AlphaFold3, while the conformations generated by AlphaFold2-Multimer using deepMSA2 alignments were better than AlphaFold3 models in terms of QS-best, ICS and IPS. These examples highlight the challenge of ranking models for some hard targets such as antibody-antigen complexes that have few good models in their model pools.

#### 3.1.2 Analysis of the prediction accuracy of MULTICOM_human in Phase 0

The average TM-score, ICS, IPS, DockQ_wave, lDDT, and QS-best of the top-1 models of MULTICOM_human for the 26 Phase 0 targets is 0.752, 0.628, 0.69, 0.584, 0.751, and 0.729, respectively. In terms of DockQ_wave, 25 out of 26 targets have top-1 models with acceptable quality (DockQ_wave >= 0.23), and 15 out of 26 targets have top-1 models with medium or high quality (DockQ_wave >= 0.49).

To further investigate how TM-score correlates with other quality metrics, we plot the TM-score of the top-1 models against their ICS, IPS, DockQ_wave, lDDT, and QS-best, respectively (Figure 5A-E). lDDT has the highest correlation with TM-score (0.77), indicating that the local structural quality score aligns well with the global structural quality score. Interface packing (IPS) has the second highest correlation (0.68) with TM-score, suggesting the packing of the interface is closely related to the global fold accuracy.

**FIGURE 5.**
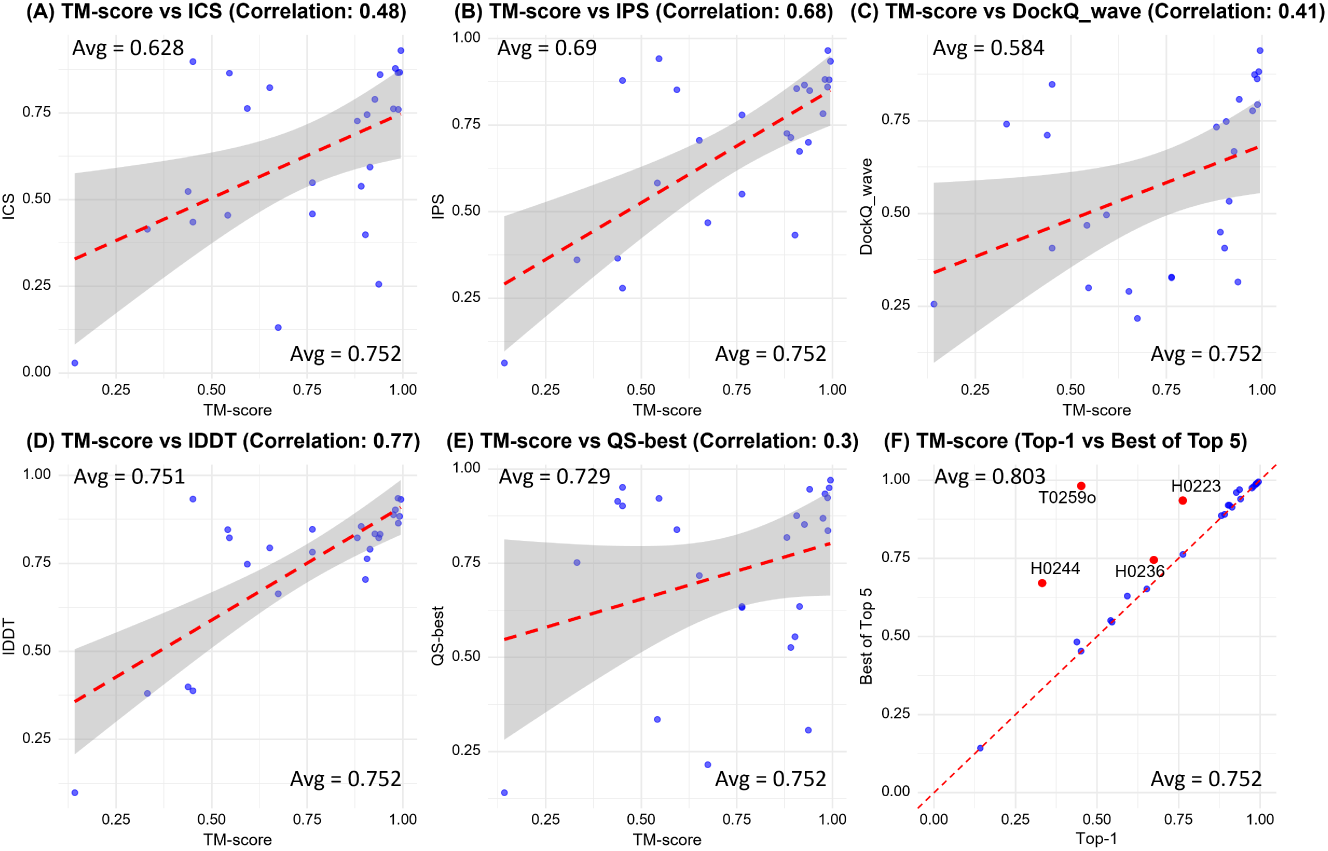
The TM-score of the top-1 model of MULTICOM_human plotted against the other five metrics of the same top-1 model and against the TM-score of the best of five models for 26 Phase 0 targets. (A) TM-Score VS ICS, (B) TM-score VS IPS, (C) TM-score VS lDDT, (D) TM-score VS DockQ_wave, (E) TM-score VS QS-best, and (F) TM-score of top-1 model VS TM-score of best of top five models. In (F), the four targets for which MULTICOM_human selected a substantially worse model than the best of top 5 models as top-1 model are highlighted in red.

In contrast, each of the three interface quality scores, ICS, DockQ_wave, QS-Best, has a moderate or low correlation of 0.48, 0.41, 0.3 with TM-score, respectively, indicating that the interface scores and the global fold quality scores are complementary and measure different aspects of structural models. A model can have a high interface score but a low global fold score and vice versa.

One such an example is T0257o (a homotrimer), a large Enterobacteria phage T5 protein that forms a long, straight, tube-like structure. However, AlphaFold3 generated bended, more globular structures, which was used by MULTI-COM_human as top-1 model. The TM-score of the model is relatively low (0.546), while its ICS is very high (0.865), indicating even though the overall fold accuracy is not high due to the bending, the local interface is still pretty accurate. This example also reveals a weakness of AlphaFold3 in predicting the structure of non-globular proteins. It also highlights the importance of using multiple complementary metrics to assess the quality of predicted structures.

Another example is T0234o (a homotrimer), a Prokaryotic polysaccharide deacetylase. The top-1 model achieves a high TM-score (0.937), indicating strong global fold accuracy. However, its DockQ_wave (0.315) and QS-Best (0.307) suggest that some local interfaces between subunits are not well predicted. This discrepancy is likely due to the predicted three-chain helical bundle in the model, different from the three-chain beta-sheet tube in the native structure. To investigate how well MULTICOM_human ranked structural models, we compare TM-score of top-1 model with that of the best of top 5 models for 26 Phase 0 targets in terms of ICS, IPS, lDDT, DockQ_wave, QS-best and TM-score (Figure 5F). The average TM-score of the best of top 5 models is 0.803, 0.051 (about 6.8%) higher than that of top-1 model (0.752). Notably, the best of top 5 models has a substantially higher TM-score than the top-1 model for four targets (H0236, H0244, T0259o, and H0223), while they have the similar score for the remaining 21 targets (81%). The results indicate that the ranking strategy of MULTICOM_human performed well for most targets but struggled with several targets.

For H0236 (correct stoichiometry: A3B6), the top five models of MULTICOM_human include AlphaFold3-generated models of both stoichiometry A3B3 and A3B6. However, MULTICOM_human incorrectly used an A3B3 model with TM-score of 0.674 as top-1 model, much worse than the A3B6 model with TM-score of 0.745 within top 5 models. Similarly, for H0244 (stoichiometry: A2B2C2), the top-1 model has the incorrectly predicted stoichiometry A1B1C1, resulting in a low TM-score (0.332), while another model with the correct stoichiometry (A2B2C2) with a much higher TM-score (0.671) was included into top five models. These two examples show that failing to select correct stoichiometry can lead to substantial loss in complex structure prediction.

For T0259o (stoichiometry: A3), the top-1 model and the best of top 5 models are very similar (TM-score = 0.971). However, when comparing them with the true structure, the top-1 model received a TM-score of 0.451, while the best of top 5 models got a TM-score of 0.982. The difference was caused by the USalign tool used to calculate TM-score, which found an optimal chain mapping between the best of top 5 models and the true structure but used a suboptimal chain mapping between the top-1 model and the true structure. This example shows that there is a need to fix the chain mapping problem in USalign to more accurately calculate TM-score for some models of homomultimers. Before the problem is fixed, if two models for homo-multimers are very similar, choose the one that likely does not have the chain mapping problem is a safer bet. In contrast, the other five metric scores for the top-1 model can be correctly calculated because they do not depend on the optimal all-chain mapping. This example underscores the importance of using multiple complementary metrics to evaluate the quality of structural models.

For H0223 (stoichiometry: A1B1C1), a hard antibody-antigen target, as discussed in the previous subsection, used a suboptimal model generated by AlphaFold2-Multimer (TM-score = 0.763) instead of the best model (TM-score = 0.935) generated by AlphaFold3 as top-1 model. This example illustrates the challenge of selecting good models for some hard antibody-antigen targets.

#### 3.1.3 Comparison of MULTICOM_human with AlphaFold2 and AlphaFold3 in Phase 0

To investigate if and how MULTICOM_human performed better than AlphaFold2-Multimer and AlphaFold3, we compare the scores of the top-1 models of MULTICOM_human against the top-1 models of AlphaFold3 (or AlphaFold2-Multimer) on the common targets which they generated structural models of the same stoichiometry for during CASP16. The top-1 models of AlphaFold3 (or AlphaFold2-Multimer) were selected by AlphaFold3 ranking scores (or AlphaFold2-Multimer confidence score) from the in-house models generated during CASP16. The comparison between MULTICOM_human and AlphaFold3 on 23 common targets is shown in Figure 6, while the comparison between MULTICOM_human and AlphaFold2-Multimer on 17 commons targets is illustrated in Figure 7.

**FIGURE 6.**
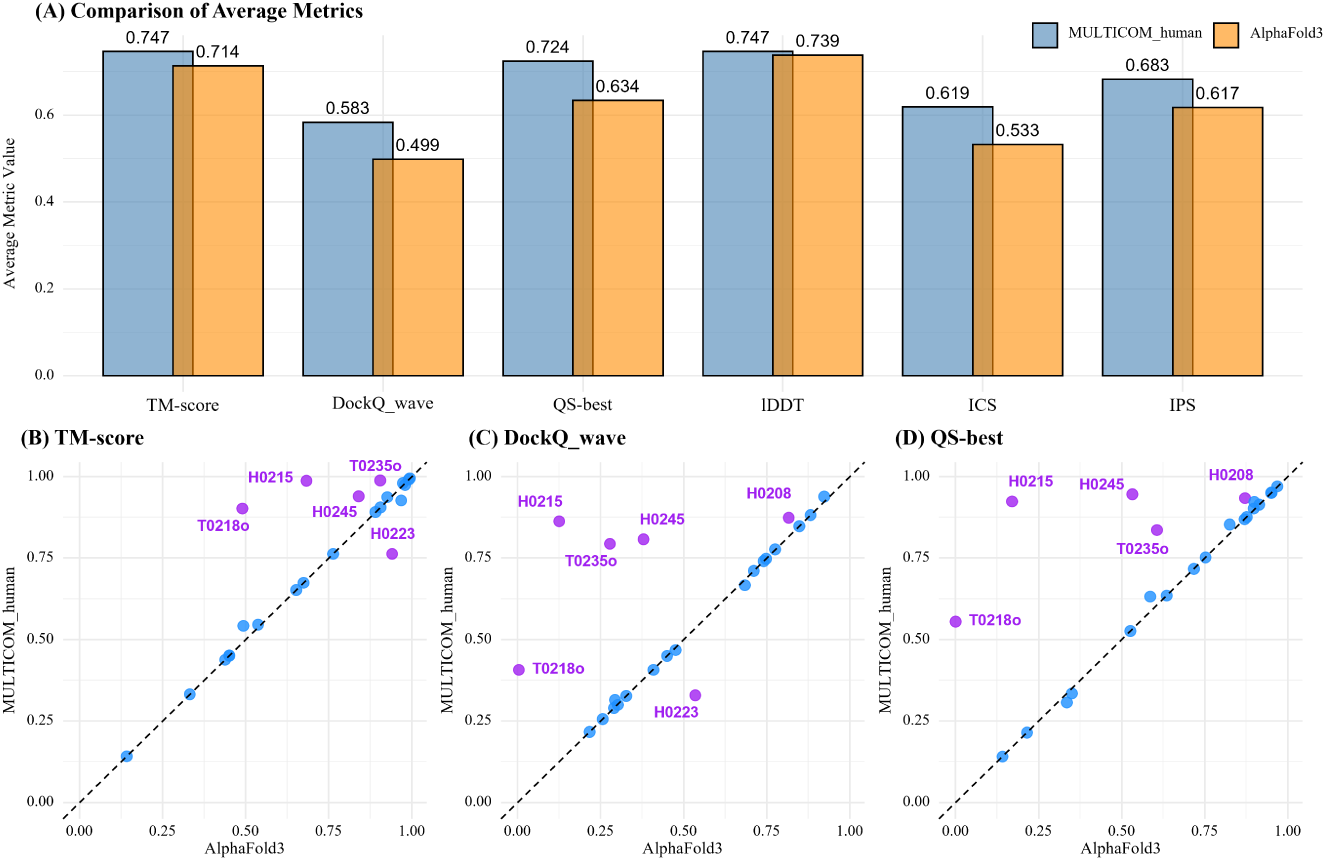
Comparison of MULTICOM_human and AlphaFold3 in Phase 0. (A) average TM-score, DockQ_wave, QS-best, lDDT, ICS and IPS of top-1 models of MULTICOM_human and AlphaFold3. The top-1 model of AlphaFold3 was selected according to AlphaFold3 ranking score. (B) Per-target TM-score of MULTICOM_human VS that of AlphaFold3. (C) Per-target DockQ_wave of MULTICOM_human VS that of AlphaFold3. (D) Per-target QS-best of MULTICOM_human VS that of AlphaFold3.

**FIGURE 7.**
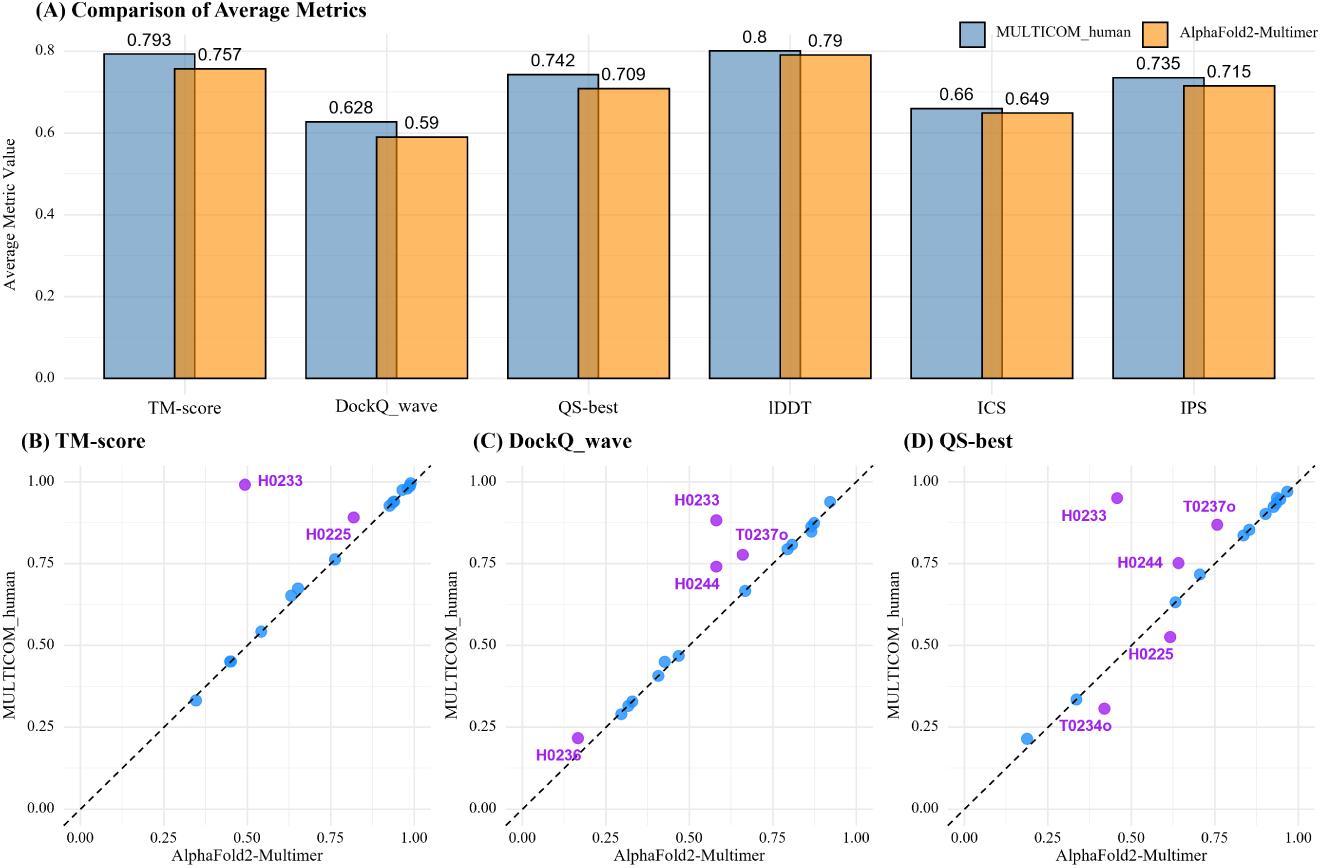
Comparison of MULTICOM_human and AlphaFold2-Multimer across key metrics in CASP16 Phase 0. (A) average TM-score, DockQ_wave, QS-best, lDDT, ICS and IPS of top-1 models for MULTICOM_human and AlphaFold2-Multimer. Per-target comparison between MULTICOM_human and AlphaFold2-Multimer in terms of (B) TM-score, (C) DockQ_wave and (D) QS-best.

On the 23 common targets, MULTICOM_human achieved higher average values than AlphaFold3 in TM-score (0.747 vs. 0.714), DockQ_wave (0.583 vs. 0.499), QS-best (0.724 vs. 0.634), lDDT (0.747 vs. 0.739), ICS (0.619 vs. 0.533), and IPS (0.683 vs. 0.617) (Figure 6A), indicating MULTICOM_human consistently performed better than AlphaFold3 on average. The improvement in terms of interface scoring metrics, including DockQ_wave, QS-best, and IPS is most pronounced, while the improvement in terms of the local quality score lDDT is marginal. This indicates that lDDT score is least sensitive to the difference in the quality of quaternary structural models because it does not assess the interaction interface or the global fold accuracy that substantially depends on the interfaces.

A detailed per-target comparison in terms of TM-score, DockQ_wave, and QS-best (Figure 6B-D) shows that MULTICOM_human performed substantially better than AlphaFold3 on five targets (T0218o, H0208, H0215, H0245, T0235o), substantially worse on one target (H0223), and similarly on the other 17 targets.

For T0218o, the better performance of MULTICOM_human is due to the exception handling of combining template-based complex models and AlphaFold3-generated tertiary structural models. Its better performance on H0208, H0215 (nanobody), H0245, and T0235o is all because MULTICOM_human used AlphaFold2-Multimer with different MSAs and parameters to generate better models than AlphaFold3. However, AlphaFold3 generated a better top-1 model for H0223 (TM-score = 0.941, DockQ_wave = 0.535), but it was not selected by MULTICOM_human as top-1 model, which was substantially better than MULTIOM_human selected top-1 model generated by AlphaFold2-Multimer (TM-score = 0.763, DockQ_wave = 0.329).

On the 17 common targets, MULTICOM_human also outperformed AlphaFold2-Multimer in TM-score (0.793 vs. 0.757), DockQ_wave (0.628 vs. 0.590), QS-best (0.742 vs. 0.709), lDDT (0.800 vs. 0.790), ICS (0.660 vs. 0.649), and IPS (0.735 vs. 0.715) (Figure 7). The reason that there are only 17 common targets is because AlphaFold2-Multimer could not generate reasonable models in a short time for several large targets on which MULTICOM_human only used AlphaFold3. The per-target comparison shows that MULTICOM_human performed much better than AlphaFold2-Multimer on two antibody targets H0233 and H0225 in terms of TM-score because it used an AlphaFold3-generated model as top-1 model that has a substantially higher TM-score than the top-ranked AlphaFold2-Multimer model. Similarly, in H0244 (a nanobody target) and T0237o, MULTICOM_human’s AlphaFold3-generated top-1 model has higher DockQ_wave and QS-best scores compared to AlphaFold2-Multimer. However, exceptions existed, such as T0234o (stoichiometry: A3), where AlphaFold2-Multimer’s top-1 model slightly outperformed MULTICOM_human’s AlphaFold3-generated model in QS-best because the former contains a correctly predicted three-chain beta-sheet tube that the latter containing a three-chain helical bundle.

Overall, the results above show that combining the strengths of AlphaFold2-Multimer with different MSAs/parameters and AlphaFold3 in MULTICOM_human in conjunction with the exception handling enabled it to perform better than both AlphaFold2-Mutimer and AlphaFold3. However, on some very hard targets (e.g., H0223), it still hard to choose better models from AlphaFold3 models and AlphaFold2-Multimer models when they differ substantially.

#### 3.1.4 Comparison of AlphaFold3 and AlphaFold2-Multimer in Phase 0

During CASP16, we observed that AlphaFold3 web server could usually generate models for large targets with thousands of residues or many chains much faster than the local AlphaFold2-Multimer. Due to the time limit, AlphaFold2-Multimer was not applied to some large targets in Phase 0. Therefore, here, we only compare them on 17 common Phase 0 targets that they both generated cohesive models during CASP16.

Figure 8 presents a comparison of the top-1 ranked models of AlphaFold3 and AlphaFold2-Multimer on 17 common targets. It is worth noting that the comparison has some limitations because more models were generally generated by AlphaFold2-Multimer than AlphaFold3 in Phase 0, and the locally installed AlphaFold2-Multimer could use diverse in-house customized MSAs and modeling parameters while we could not use them with AlphaFold3 web server.

**FIGURE 8.**
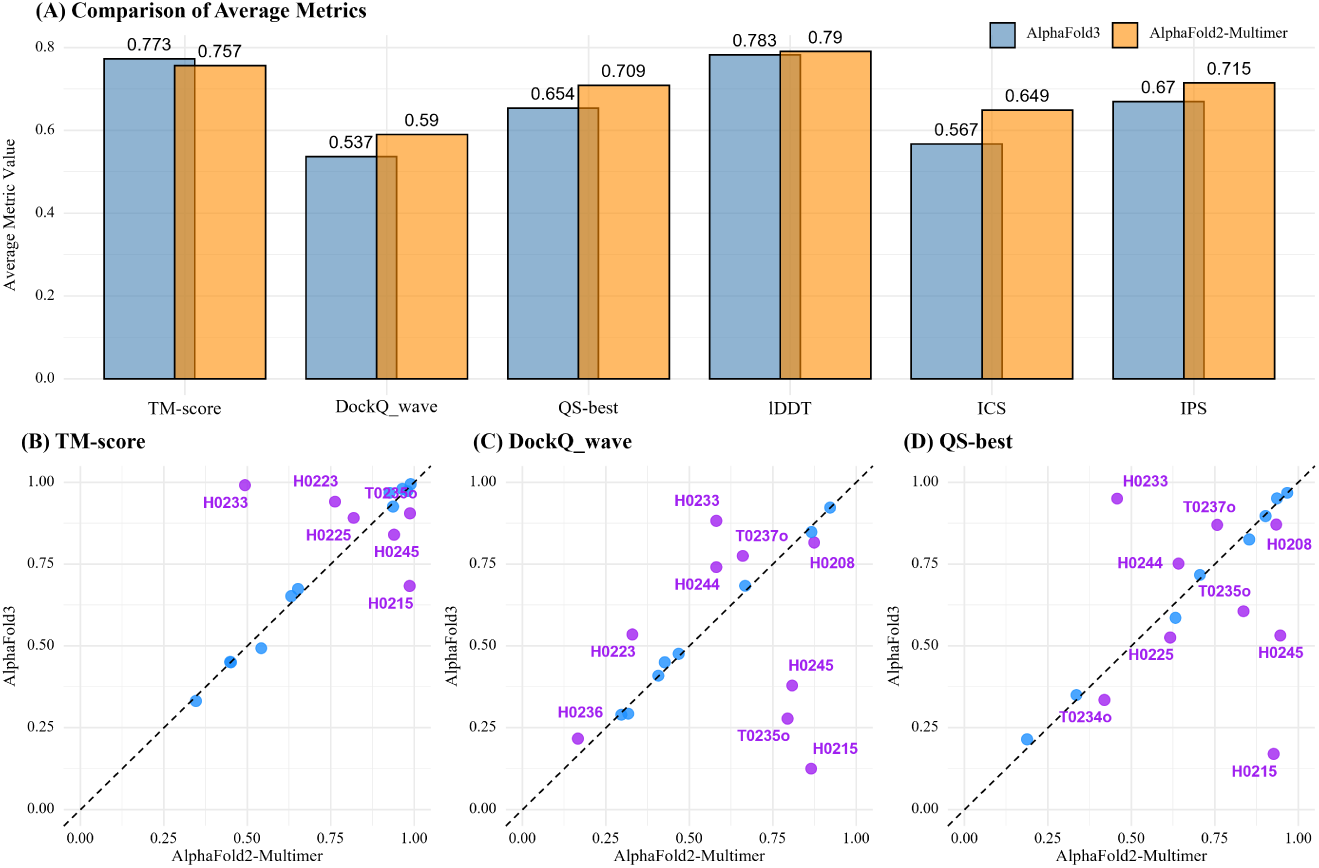
Comparison of AlphaFold3 and AlphaFold2-Multimer on 17 common targets in Phase 0. (A) average TM-score, DockQ_wave, QS-best, lDDT, ICS and IPS for top-1 ranked models of AlphaFold3 and AlphaFold2-Multimer. Per-target comparison between AlphaFold3 and AlphaFold2-Multimer in terms of (B) TM-score, (C) DockQ_wave and (D) QS-best.

Overall, on the 17 common targets, AlphaFold3 demonstrated a slight advantage in global structural accuracy, with an average TM-score of 0.773, higher than 0.757 of AlphaFold2-Multimer. However, AlphaFold2-Multimer out-performed in interface-specific metrics, achieving higher DockQ_wave (0.59 vs. 0.537), QS-best (0.709 vs. 0.654), ICS (0.649 vs. 0.567), and IPS (0.715 vs. 0.67). Both methods had comparable lDDT scores, with AlphaFold2-Multimer slightly ahead (0.790 vs. 0.783), indicating similar local structural quality.

A per-target comparison reveals that AlphaFold3 excels in antibody targets (Figure 8B-D), achieving higher TM-scores in H0223 (0.941), H0225 (0.891), and H0233 (0.991) compared to AlphaFold2-Multimer (0.763, 0.819, 0.493) (Figure 9A-C. However, AlphaFold2-Multimer also demonstrated superior interface accuracy in certain cases. In H0215 (mNeonGreen with bound nanobody), AlphaFold2-Multimer achieved substantially higher TM-score (0.987 vs. 0.683), DockQ_wave (0.865 vs. 0.125), and QS-best (0.926 vs. 0.170) by using multimer_v1 weights to gener-ate 1,000 models (Figure 9D). For H0245 with a shallow MSA, AlphaFold2-Multimer achieved a higher TM-score (0.94 vs. 0.84) and significantly better interface metrics, with DockQ_wave (0.808 vs. 0.379) and QS-best (0.946 vs. 0.532) than AlphaFold3 because it used the customized FoldSeek structure-based MSA. The superior performance of AlphaFold2-Multimer on this target is likely due to the improved MSA input rather than its neural network model. For homomultimer T0235o (stoichiometry: A6), AlphaFold2-Multimer achieved much higher DockQ_wave (0.794 vs. 0.278), QS-best (0.836 vs. 0.606), and TM-score (0.988 vs. 0.905) than AlphaFold3.

**FIGURE 9.**
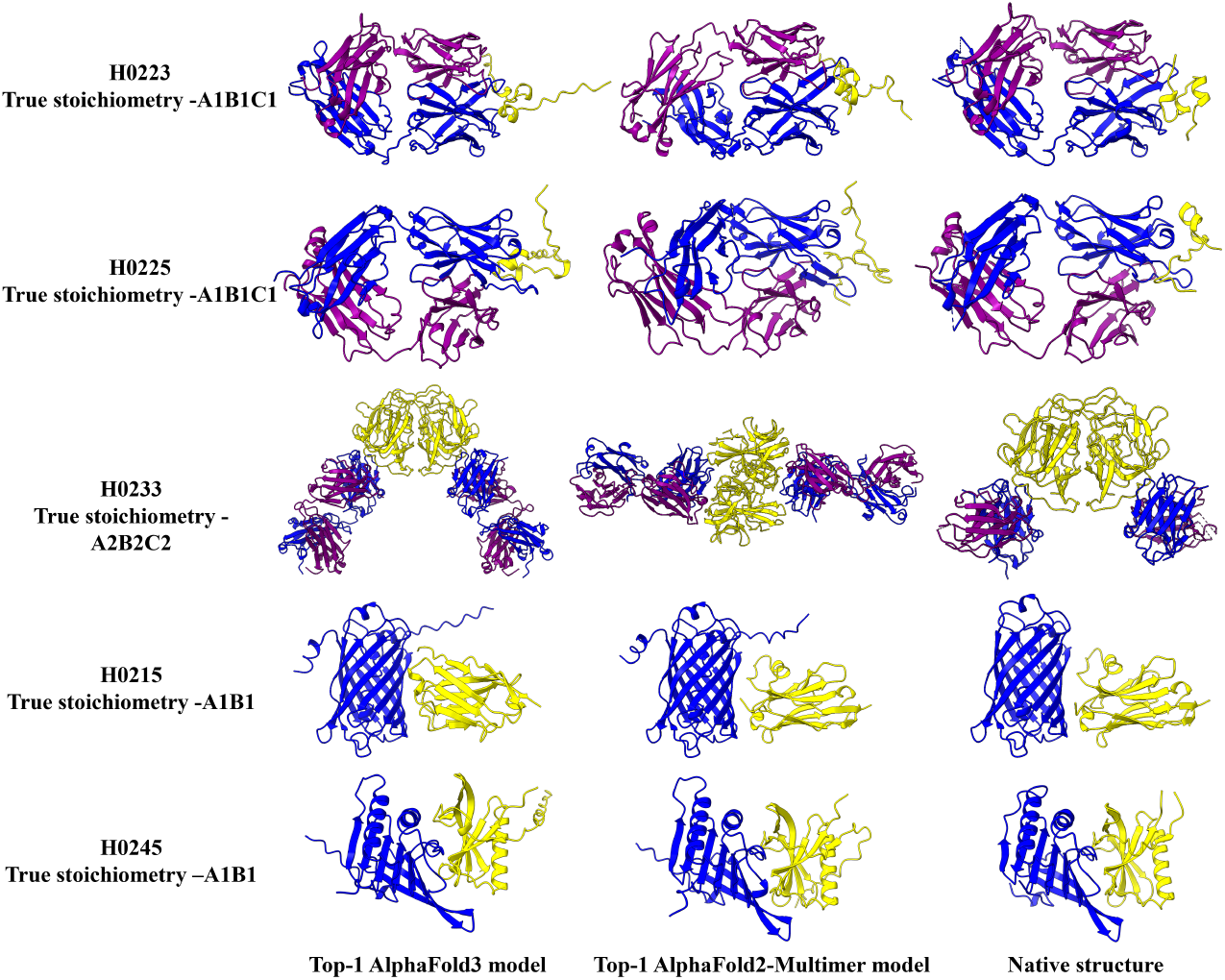
Structure comparison of AlphaFold3 and AlphaFold2-Multimer on the antibody targets (H0223, H0225, H0233), an nanobody target (H0215) and a FUNComplex target H0245. Left: top-1 AlphaFold3 model; Middle: top-1 AlphaFold2-Multimer model; Right: Native structure

The results above underscore the complementary strengths of AlphaFold3 and AlphaFold2-Multimer and the importance of using diverse MSAs / modeling parameters with AlphaFold2-Multimer to increase the likelihood of generating better models for hard targets. We expect that using diverse MSAs and modeling parameters can also improve the accuracy of AlphaFold3, even though we were not able to do it during CASP16.

### 3.2 Complex Prediction Results with Stoichiometry Information in Phase 1

#### 3.2.1 Comparison of MULTICOM Predictors with other CASP16 predictors in Phase 1

Figure 10A visualizes the cumulative Z-scores of the top-1 models of top 20 out of 88 CASP16 predictors on 39 multimeric targets with native structures available. All the MULTICOM predictors ranked among top-15 predictors. Particularly, MULTICOM_human ranked 4th, right after three predictors from Yang Group. All the MULTIOCM predictors substantially outperformed the standard AlphaFold3 predictor (AF3-server run by Dr. Eloffson’s group) that was ranked 14th.

**FIGURE 10.**
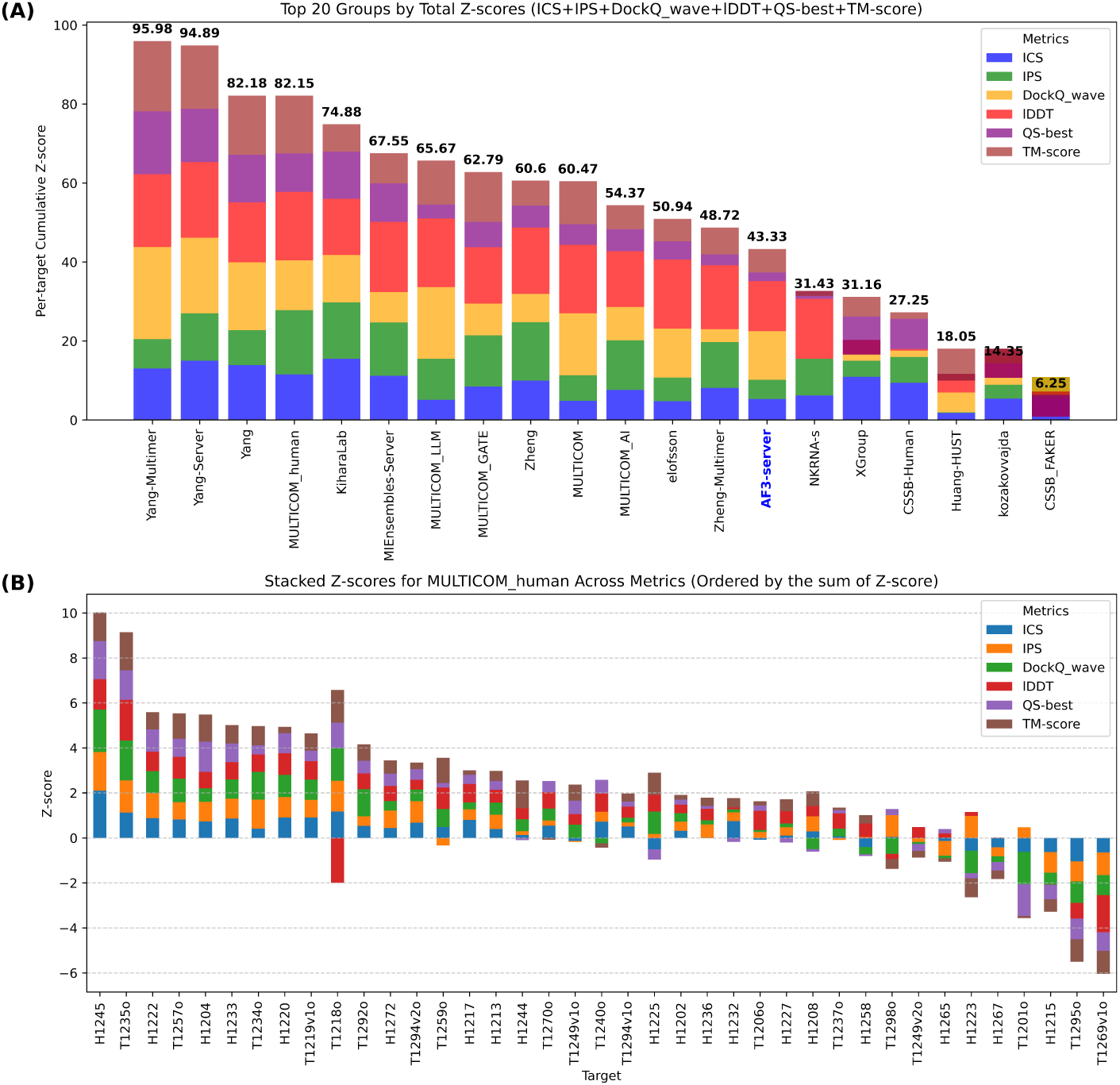
The performance of top 20 out of 88 predictors in Phase 1. (A) the cumulative Z-scores of top 20 predictors across six metrics for their top-1 models for 39 targets. The standard AlphaFold3 predictor (AF3-server) is highlighted in blue font, which is ranked at 14th place with the cumulative Z-score 43.33; (B) The per-target Z-score of top-1 model of MULTICOM_human.

Figure 10B illustrates the per-target Z-score of top-1 models of MULTICOM_human. It performed better than average on 30 out of 39 targets (cumulative Z-score > 0), such as H1245 and T1235o, as it did for most targets in Phase 0, such as H0245 and T0235o (the counterparts of H1245 and T1235o) for similar reasons.

Despite its strong performance on most targets, MULTICOM_human performed worse than an average predictor on 9 out of 39 targets (Z-score < 0), which are described below.

For T1298o (a homodimer), whose subunit (chain) has two domains linked by a flexible linker, MULTICOM_human selected a model with a straight linker generated by AlphaFold2-Multimer as top-1 model that has few interactions between the two domains with each subunit. It is substantially worse than the models with a bended linker generated by AlphaFold3 that allows for more interactions between the two domains. This example demonstrates the challenge of selecting better models for targets with flexible domain linkers when the models generated by AlphaFold2-Multimer and AlphaFold3 differ a lot.

For T1249v2o, which shares the same sequence as T1249v1o but adopts a more closed conformation, AlphaFold3 correctly predicted both open and closed conformations with high accuracy. However, MULTICOM_human failed to select the correct conformation for this target, resulting in suboptimal prediction, underscoring the challenge of model selection for structurally dynamic targets that may have multiple conformations.

For H1265 (stoichiometry: A9B18), a large filament target consisting of 27 chains in total, MULTICOM_human chose a cyclic conformation incorrectly predicted by AlphaFold3 as top-1 model. Notably, no in-house models were generated with the correct fold (i.e., TM-score > 0.5) or medium/high-quality interfaces (i.e., DockQ_wave > 0.49) for this target. H1265 stands out as one of the most challenging cases in CASP16, with no participating group successfully producing a high-accuracy model for it.

For H1223, an antibody target, AlphaFold2-Multimer and AlphaFold3 produced very different conformations, making it difficult to determine which was better. The top-1 model selected by MULTICOM_human, generated using AlphaFold2-Multimer, had a lower TM-score (0.763) and DockQ_wave (0.329) compared to a much better AlphaFold3 model (TM-score = 0.941, DockQ_wave = 0.546). This case highlights the challenge of selecting good models for antibody-antigen targets, particularly when AlphaFold3 and AlphaFold2-MUltimer do not agree with each other.

For H1267 (stoichiometry: A2B2, tetramer), the challenge is to determine the interaction interface between two dimers (A1B1) in the tetramer, for which AlphaFold3 and AlphaFold2-Multimer made different predictions. The top-1 model selected by the default ranking in MULTICOM_human was generated by AlphaFold2-Multimer with DeepMSA2 alignments, which has a better TM-score (0.847), DockQ_wave (0.362) and QS-best (0.723) compared with the top-1 AlphaFold3 model (TM-score: 0.453, DockQ_wave: 0.286, QS-best: 0.551). However, a manual adjustment was made to select the AlphaFold3 model as top-1 model of MULTICOM_human. This example signifies that manual model ranking does not always work better than automated model ranking.

For T1201o (a homodimer), an early-stage target, where AlphaFold3 was just released and was used to generate only 10 models for it, AlphaFold2-Multimer produced models with high TM-scores around 0.9 but varying interface quality (ICS), making ranking the models difficult. MULTICOM_human selected a sub-optimal model with low interface quality. This example highlights the need of better QA methods that account for multimeric interface interactions rather than relying solely on global structural accuracy.

For H1215 (stoichiometry: A1B1), mNeonGreen with bound nanobody, is a hard target whose models have various interaction interfaces between mNeonGreen and nanobody, making it hard to choose a correct one. MULTI-COM_human chose a low-quality model with the incorrect interface as top-1 model in Phase 1, even though it chose a high-quality model for the same target (H0215) with a TM-score of 0.987 and DockQ_wave of 0.863 as top-1 model in Phase 0. This example demonstrates that generating more models in Phase 1 did not always improve prediction over Phase 0 because it could make model ranking more complicated by adding highly scored false positive models.

For T1295o (stoichiometry: A8), a nanoparticle, MULTICOM predictors generated multiple conformations using both AlphaFold3 and AlphaFold2-Multimer. Even though some models generated by AlphaFold2-Multimer with DeepMSA2 alignments were of better quality, MULTICOM_human failed to select them as top-1 model. This example reinforces the observation that it is challenging to select models from many different conformations are generated by AlphaFold2-Multimer and AlphaFold3. This target is also one of the hardest targets in CASP16 that no predictor was able to generate models of correct global fold, even though some of CASP16 predictors predicted some interaction interfaces correctly.

Finally, for T1269v1o, a filament target, MULTICOM predictors used the template-based modeling to build models from a template (PDB code: 7A5O) with 100% sequence identity to the target. However, the native structure adopted a very different conformation, which could have been correctly predicted by AlphaFold3. This over-reliance on sequence-identical templates led to incorrect structural predictions. It also highlighted the difficulty of predicting or selecting structures for targets that may adopt multiple different conformations.

In summary, MULTICOM predictors performed well on most Phase 1 targets, but the model ranking strategy for antibody/nanobody-related proteins, filamentous structures, and complexes with flexible or multiple conformations needs to be improved. The model generation for very large complexes or non-globular filaments also needs improvement.

#### 3.2.2 Comparison between MULTICOM_human, AlphaFold3 and AlphaFold2-Multimer in Phase 1

The average TM-score, ICS, IPS, DockQ_wave, lDDT, and QS-best of the top-1 models of MULTICOM_human for the 39 Phase 1 targets is 0.797, 0.622, 0.69, 0.558, 0.817, and 0.677, respectively. In terms of DockQ_wave, 34 out of 39 targets (87%) have top-1 models with acceptable quality (DockQ_wave >= 0.23), and 24 out of 39 targets (62%) have top-1 models with medium or high quality (DockQ_wave >= 0.49).

Figures 11 and 12 compare the top-1 models of MULTICOM_human with those of AlphaFold3 and AlphaFold2-Multimer on the common targets in terms of TM-score, DockQ_wave, QS-best, lDDT, ICS, and IPS of their top-1 ranked models, respectively. Similarly as in Phase 0, on the 31 common Phase 1 targets for which AlphaFold3 models were generated, MULTICOM_human achieved consistently higher average scores in terms of all the metrics: TM-score (0.789 vs. 0.769), DockQ_wave (0.590 vs. 0.588), QS-best (0.697 vs. 0.688), lDDT (0.833 vs. 0.829), ICS (0.654 vs. 0.641), and IPS (0.724 vs. 0.699).

**FIGURE 11.**
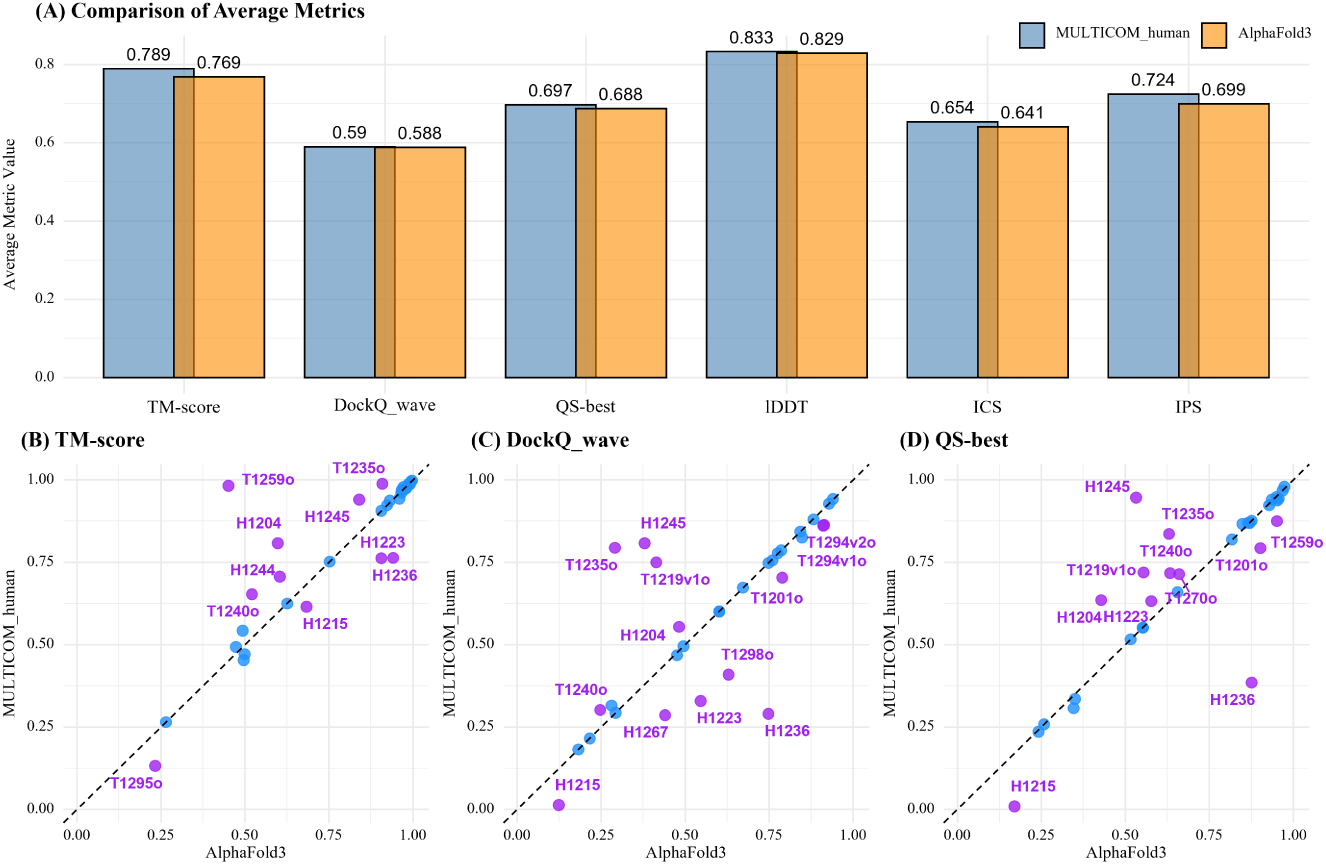
Comparison of MULTICOM_human and AlphaFold3 across key metrics in terms of top-1 models for 31 common Phase 1 targets. The top-1 models for AlphaFold3 were selected based on the ranking scores. (A) average TM-score, DockQ_wave, QS-best, lDDT, ICS and IPS for MULTICOM_human and AlphaFold3; Per-target comparison of (B) TM-score, (C) DockQ_wave and (D) QS-best.

**FIGURE 12.**
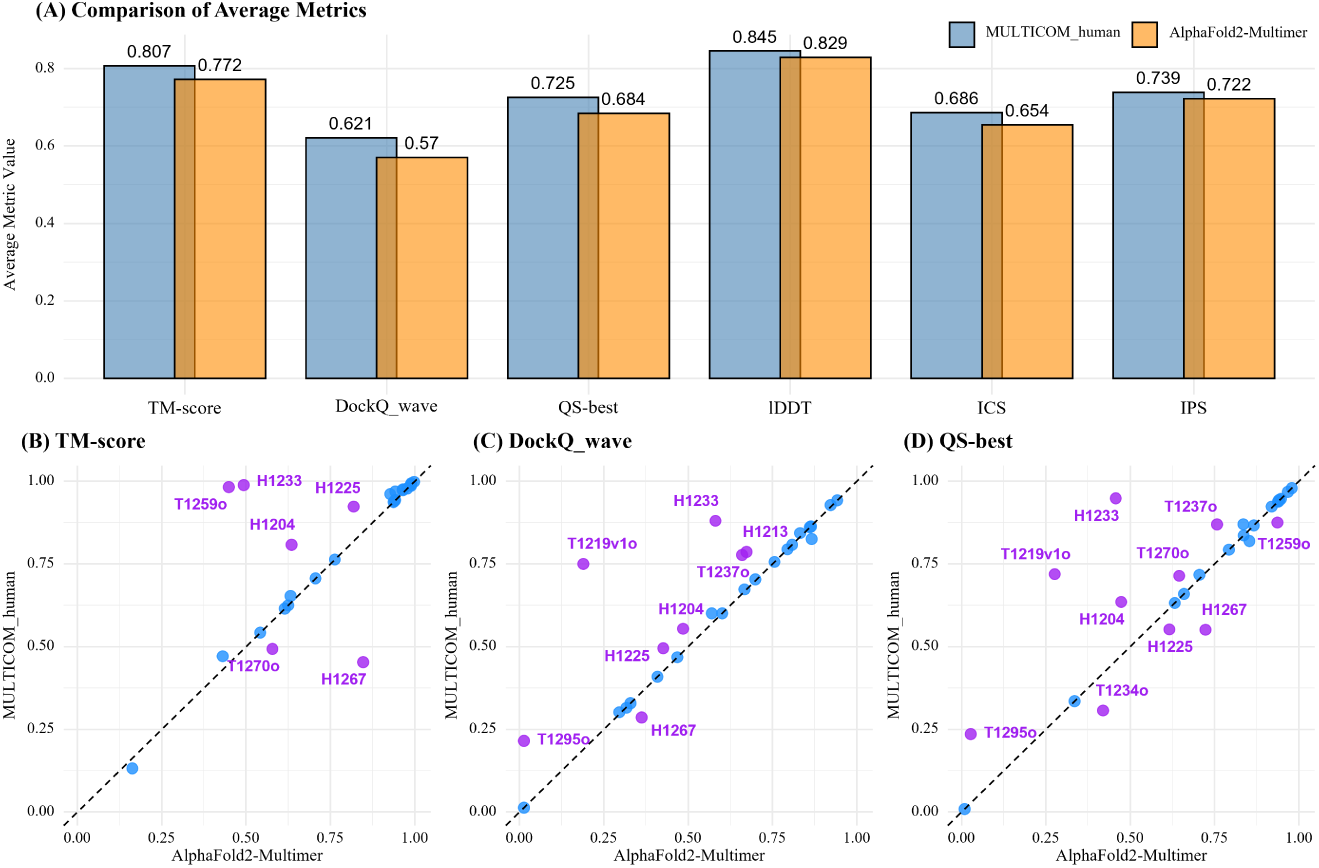
Comparison of MULTICOM_human and AlphaFold2-Multimer across key metrics in terms of top-1 models for 27 common Phase 1 targets. The top-1 models for AphaFold2-Multimer were selected based on the confidence scores. (A) average TM-score, DockQ_wave, QS-best, lDDT, ICS and IPS for MULTICOM_human and AlphaFold2-Multimer; Per-target comparison of (B) TM-score, (C) DockQ_wave and (D) QS-best.

According to the per-target comparison in Figure 11B, MULTICOM_human substantially outperformed AlphaFold3 on several homo-multimers. For example, on T1235o(TM-score: 0.988 vs. 0.909; DockQ_wave: 0.794 vs. 0.291) and T1240o (TM-score: 0.653 vs. 0.521; DockQ_wave: 0.302 vs. 0.247), MULTICOM_human achieved better results because it used an AlphaFold2-Multimer generated model that is better than AlphaFold3 models as top-1 model. For T1259o (TM-score: 0.982 vs. 0.451; DockQ_wave: 0.825 vs. 0.848), MULTICOM_human used an AlphaFold3 model with a higher AlphaFold2-Multimer confidence score and that did not have the chain mapping issue with most of other models through manual ranking adjustment, leading to better overall performance compared to the top-1 AlphaFold3 model ranked by AlphaFold3 ranking score. However, the manually selected AlphaFold3-generated top-1 model of MULTICOM_human for T1295o (TM-score: 0.132 vs 0.233) was worse than the top-ranked AlphaFold3 model of different conformation that has the same AlphaFold3 ranking score. These two examples show that the manual ranking adjustment yielded inconsistent results.

For hetero-multimers, MULTICOM_human substantially outperformed AlphaFold3 on several targets such as H1245 and H1204 due to either better top-1 model selection or using AlphaFold2-Multimer models generated with customized MSAs or diverse modeling parameters, while it also substantially performed worse than AlphaFold3 due to failed top-1 model selection on several targets, such as H1215 (a nanobody target), H1223 (an antibody target), Similarly as in Phase 0, on the 27 common Phase 1 targets, MULTICOM_human consistently outperformed AlphaFold2-Multimer across all metrics, achieving higher TM-score (0.807 vs. 0.772), DockQ_wave (0.621 vs. 0.570), QS-best (0.725 vs. 0.684), lDDT (0.845 vs. 0.829), ICS (0.686 vs. 0.654), and IPS (0.739 vs. 0.722).

Figure 12B highlights target-specific differences between MULTICOM_human and AlphaFold2-Multimer. MUL-TICOM_human substantiallly outperformed AlphaFold2-Multimer on several targets, such as H1233, H1204, H1213, T1219v1o, and T1295o in terms of TM-score and/or the interface metrics due to the use of AlphaFold3 models or better model ranking. For T1259o, the top-1 model of MULTICOM_human achieves a TM-score of 0.982 compared to AlphaFold2-Multimer’s 0.449 because USalign could find an optimal chain mapping between the former and the experimental structure but not between the latter and the experimental structure. However, MULTICOM_human’s QS-best (0.875) is not much lower than AlphaFold2-Multimer’s 0.936, suggesting the interface quality metric is not sensitive to the chain mapping.

For one target, H1267 (stoichiometry: A2B2), MULTICOM_human substantially underperformed AlphaFold2-Multimer in terms of the three metrics: TM-score (0.453 vs. 0.847), DockQ_wave (0.286 vs. 0.362), and QS-best (0.551 vs. 0.723). Although the AlphaFold2-Multimer model was ranked 1st in MULTICOM_human’s default ranking, a manual adjustment was made to use an AlphaFold3 model with much lower quality as top-1 model.

Similarly as in Phase 0, the Phase 1 results above demonstrate that using both AlphaFold3 and AlphaFold2-Multimer with diverse MSAs and modeling parameters to generate structural model as well as multiple model ranking strategies improves complex structure prediction accuracy over either AlphaFold3 or AlphaFold2-Multimer consistently.

Moreover, as in Phase 0, we conducted a head-to-head comparison of top-1 models of AlphaFold2-Multimer and AlphaFold3 on 27 common targets, excluding some very large targets that only AlphaFold3 could generate structures in a short time and some early targets that AlphaFold3 was not applied to (Figure 13).

**FIGURE 13.**
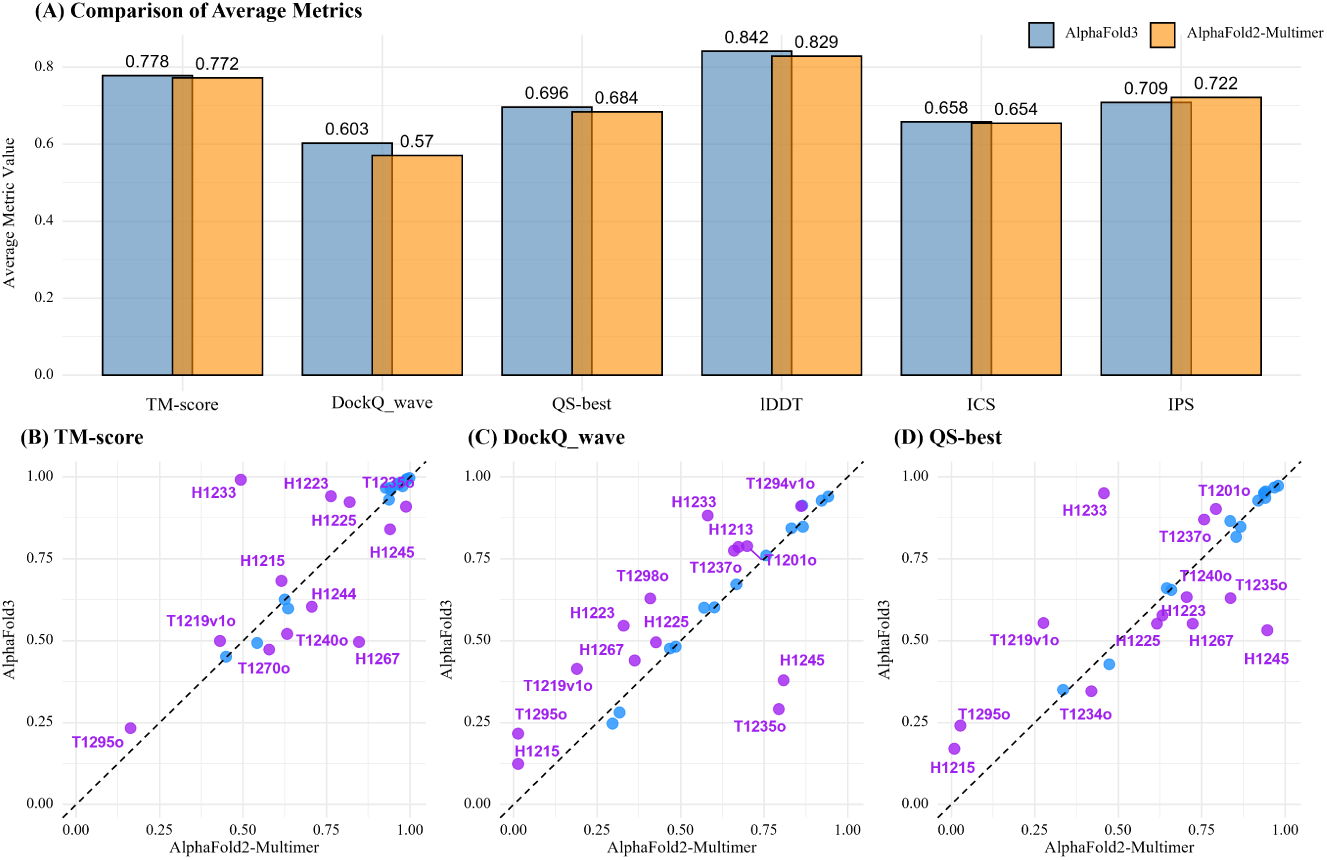
Comparison of AlphaFold3 and AlphaFold2-Multimer across key metrics in terms of top-1 models in CASP16 Phase 1.(A) average TM-score, DockQ_wave, QS-best, lDDT, ICS and IPS for AlphaFold3 and AlphaFold2-Multimer; Per-target comparison of (B) TM-score, (C) DockQ_wave and (D) QS-best.

AlphaFold3 performed slightly better than AlphaFold2-Multimer in terms of average TM-score (0.778 vs. 0.772), DockQ_wave (0.603 vs. 0.570) and QS-best (0.696 vs. 0.684), ICS values (0.658 vs. 0.654), lDDT scores (0.842 vs. 0.829), but slightly worse in terms of IPS score (0.709 vs. 0.722). The comparison result here is consistent with the comparison between the two in Phase 0 in terms of global fold accuracy (TM-score), but is somewhat different from Phase 0 in terms of interface and local quality metrics (e.g. DockQ_wave, QS-best, ICS, lDDT). The difference is likely due to the fact that there are more targets and an increased number of structural models generated by AlphaFold3 in Phase 1.

Because more targets used in Phase 1 than in Phase 0 can possibly lead to a more reliable estimate and AlphaFold2-Multimer benefited from our in-house customized MSAs, it is reasonable to conclude that AlphaFold3 performs slightly better than AlphaFold2-Multimer on average. However, they are also complementary, i.e., each of them may work better on some specific targets.

The scatter plots in Figure 13 provide a per-target comparison between AlphaFold3 and AlphaFold2-Multimer. AlphaFold3 substantially outperformed AlphaFold2-Multimer on several targets, such as three antibody targets (H1223, H1225, H1233) as in Phase 0 in terms of most evaluation metrics, particularly TM-score, while it significantly underperformed AlphaFold2-Multimer on several targets, such as T1235o, H1245, and some nanobody targets (e.g., H1204 and H1244). For another nanobody target H1215, AlphaFold2-Multimer also generated a high-quality model (e.g., the best model of H0215, TM-score: 0.987) in Phase 0, but its confidence score was slightly lower than a low-quality model generated by AlphaFold2-Multimer in Phase 1 and therefore was not selected as top-1 model. This example reveals a side effect of generating many models in some cases, introducing some high-scoring false positives.

Overall, there is a strong complementarity between AlphaFold3 and AlphaFold2-Multimer. Using both of them to generate models for model selection in MULTICOM_human can improve prediction accuracy.

## 4 DISCUSSION

MULTICOM predictors built on the MULTICOM4 system delivered an outstanding performance in both Phase 0 without stoichiometry information and in Phase 1 with stoichiometry information in comparison with other CASP16 predictors. Its advantage mainly stemmed from using both AlphaFold2 and AlphaFold3 to generate many structural models, using diverse MSAs and modeling parameters with local AlphaFold2-Multimer, applying multiple complementary QA methods to rank and select models, better stoichiometry prediction in Phase 0, and handling exceptional targets (e.g., extremely large ones) that AlphaFold could not deal with. MULTICOM predictors, such as MULTICOM_human, also outperformed both AlphaFold3 and AlphaFold2-Multimer executed in the MULTICOM4 system by integrating predictions from both methods and refining model selection beyond their default ranking scores.

Among several individual QA methods used in MULTICOM4, the average of GATE and the AlphaFold2-Multimer confidence score as well as GATE was generally more effective than AlphaFold2-Multimer confidence score or the pairwise structural similarity score (PSS), considering their performance in both Phases 0 and 1 (data not shown). The AlphaFold3 ranking score is also useful, but can only be applied to AlphaFold3-generated models. Overall, using multiple QA methods and including alternative conformations into top five models via removing highly similar models or model clustering for hard targets can include some good model into top 5 models rather effectively. However, the hybrid model ranking strategy still could not consistently select good models as top-1 model for all hard targets with few good models generated, particularly when multiple different, but highly-scored conformations were generated by AlphaFold2 and AlphaFold3 for some targets (e.g., H1225 and H1223). Therefore, model quality assessment and ranking is a major challenge in improving protein complex structure prediction.

AlphaFold3 and AlphaFold2-Multimer exhibited distinct but complementary strengths in multimeric structure prediction. AlphaFold3 generally performed better on most antibody-related targets such as H0223 and H0225, where it achieved higher TM-scores and captured overall fold accuracy more reliably. In contrast, AlphaFold2-Multimer empowered by diverse MSAs and different modeling parameters may excel on some targets with shallow MSAs (H0245/H1245) and nanobody targets (e.g., H0215) as well as in interface modeling for some targets (e.g., T0235o). However, the advantage of AlphaFold2-Multimer on some targets over AlphaFold3 such as H0245 may be due to better customized MSAs rather than its own better prediction capability. Overall, AlphaFold3 may be a little more accurate than AlphaFold2-Multimer on average.

It is also important to use both global fold and interface quality metrics to comprehensively evaluate the quality of structural models for comparing the performance of different predictors. For example, for H1223, AlphaFold3 produced a more globally accurate structure in terms of TM-score, but AlphaFold2-Multimer generated models with slightly better interface contacts in terms of the interface metrics. By using both metrics, their strengths and weaknesses can be objectively assessed.

Finally, in comparison with CASP15 results of MULTICOM3, the model generation capability of MULTICOM4 has been substantially improved largely due to the incorporation of AlphaFold3, particularly for large protein complexes and antibody targets. During CASP16, MULTICOM4 was able to generate at least one model with correct fold (TM-score > 0.5) for 36 out of 39 targets in Phase 1. It is worth noting that the commonly used threshold for determining if the fold of a tertiary structure is correct is sometime too low for quaternary structures of small complexes with few subunits such as dimers. For all antibody and nanobody targets, it also generated at least one model with above-medium interface quality (e.g., DockQ_wave >= 0.49). But it failed to generate a model with correct fold for 3 large filament targets, including T1219v1o (stoichiometry: A8), H1265 (stoichiometry: A9B18) and T1295o (stoichiometry: A8). The difficulty for AlphaFold 2 and 3 to generate correct structures for non-globular filaments is probably due to the fact that its training data contain mostly globular structures. Improving structure generation for large non-globular proteins is an important direction to pursue in the future.

## 5 CONCLUSION

We developed the MULTICOM4 system for protein complex structure prediction and blindly tested it in CASP16 as several MULTICOM predictors. The MULTICOM predictors ranked among the top predictors in protein complex structure prediction with / without stoichiometry information. The experiment demonstrates that applying both AlphaFold2 and 3 for model generation, using diverse MSAs and modeling parameters with AlphaFold, combining multiple QA methods to rank models, using accurate stoichiometry prediction, and applying exception handling to deal with the limitation of AlphaFold can systematically improve AlphaFold2 and 3-based protein complex structure prediction. Moreover, the experiment identified two key challenges to further improve protein complex structure prediction, including accurately ranking good models at top for hard targets such as antibodies and generating good models for non-globular large protein complexes such as filament.

## Author contributions

Jian Liu: Methodology; investigation; writing - original draft, reviewing and editing; software. Pawan Neupane: Investigation; writing - reviewing and editing. Jianlin Cheng: Conceptualization; methodology; investigation; writing-original draft, reviewing and editing.

## Acknowledgments

This work was supported by a U.S. NSF grant (DBI2308699) and U.S. NIH grants (R01GM093123 and R01GM146340) awarded to JC. We thank CASP16 organizers and assessors for sharing CASP16 data and results.

## Availability statement

The source code of MULTICOM4 is available at https://github.com/BioinfoMachineLearning/MULTICOM4. Addi-tionally, the data used in this study are available on the CASP website at https://predictioncenter.org/casp16/ results.cgi?tr_type=multimer.

